# Behavioral variation affects persistence of an experimental food-chain

**DOI:** 10.1101/2025.07.04.663144

**Authors:** Pragya Singh, Gaurav Baruah, Caroline Müller

## Abstract

Intraspecific behavioral variation in prey could alter predator–prey interactions, yet its effects on temporal dynamics and food-web persistence remain underexplored. Pea aphids (*Acyrthosiphon pisum*) exhibit dropping behavior in response to predators like the seven-spot ladybird (*Coccinella septempunctata*). This response could be an effective anti-predator defense but could be costly in terms of energy expenditure and time not available for feeding. To investigate the impact of behavioral variation on food-chain persistence and dynamics, we used a tri-trophic experimental system with *Vicia faba* (plant), pea aphids (prey), and seven-spot ladybirds (predator), implementing three aphid behavioral treatments: droppers, non-droppers, and a mix of both droppers and non-droppers. To minimize genetic differences, we used clonal aphid populations across all treatments. We then tracked predator-prey population dynamics and species persistence over 25 days. Our results showed that aphid dropping behavior reduced food-chain persistence, with extinction risk significantly higher in dropper treatments than in the mix or non-dropper treatments. Ladybirds persisted across treatments, although they showed a steeper decline in abundance in the dropper treatment. In the mixed behavioral treatment, they had an intermediate persistence, suggesting a buffering effect of behavioral variation. Trophic food-chain state transitions also differed by treatment, with tri-trophic states most stable in the non-dropper, and least frequent in the dropper treatment. Furthermore, our results showed a trend of dropper treatments becoming more stable and robust towards the end of the experiment. These results demonstrate that prey behavior influences the persistence and dynamics of food-chains, with important implications for behavior-driven community dynamics.

## 1. Introduction

Individual variation is ubiquitous in nature, and has been suggested to impact ecosystem function diversity (Barbour *et al*. 2016; Baruah 2022; Bolnick *et al*. 2011; Uriarte & Menge 2018) and population persistence (Baruah *et al*. 2021; Gibert 2016). Such variation can arise due to genetic variation as well as due to variation caused by the environment. Theoretical studies have consistently suggested the importance of such variation in species on maintenance of diversity and ecosystem functions (Barabas & D’Andrea 2016; Baruah *et al*. 2025; Hart *et al*. 2016), but the focus has mainly been on the genetic causes of such variation. The importance of genetic variation within species for ecosystem functioning and dynamics is well established, however, the ecological effects of intraspecific behavioral variation remain relatively understudied. Behavioral variation in traits such as boldness, activity level, and escape responses have been observed in empirical studies across diverse taxa (Laskowski *et al*. 2022; Magnhagen & Staffan 2005; Sih *et al*. 2012; Singh *et al*. 2023). Such intraspecific behavioral variation can alter predation risk and interaction outcomes of species persistence (Foster *et al*. 2017; McGhee *et al*. 2013; Réale & Festa-Bianchet 2003). For example, bold or highly active individuals may experience higher encounter rates with predators (Belgrad & Griffen 2016; Magnhagen & Staffan 2005; Singh *et al*. 2023). These behavior-mediated differences can shape not only individual survival but also species abundances and interaction strengths within communities. Such variation may consequently scale up and impact community stability and persistence.

While these studies highlight how prey behavior influences short-term predator-prey interactions, such as who gets eaten or/and how fast, a critical gap remains in our understanding of how behavioral variation impacts the temporal dynamics of species as well as community persistence. Few studies have addressed whether intraspecific behavioral diversity contributes to the persistence or temporal stability of predator-prey systems (Barbour *et al*. 2019). Yet, ecological theory suggests that trait variation can stabilize populations and also buffer populations against extinction (Barabas & D’Andrea 2016; Baruah & Lakämper 2024), potentially stabilizing dynamics over time. These individual-level differences can scale up by altering prey population growth, modifying species interactions (Barbour *et al*. 2022; Baruah & Wittmann 2024; Gibert & Brassil 2014), and changing the strength of trophic cascades (DeLong *et al*. 2015). For instance, if predators preferentially remove herbivores with certain behaviors, plants may experience different grazing pressure. As a result, prey behavioral diversity adds complexity to food webs that can stabilize or destabilize interactions and impact biodiversity (Schmitz *et al*. 1997).

Intraspecific variation in behavioural traits is widespread and can arise from genetic differences, state-dependent feedbacks, life-history trade-offs, fluctuating selection, social interactions, or developmental noise, each driven by distinct ecological and evolutionary pressures (Barbour *et al*. 2022; Dochtermann & Royauté 2019; Laskowski *et al*. 2022; Réale *et al*. 2010). For example, animals often exhibit consistent individual differences in boldness or activity levels, which can manifest as behavioural syndromes, i.e., suits of correlated behaviours expressed across contexts (Sih *et al*. 2012). Such syndromes may limit behavioral plasticity; for instance, more active individuals may maintain high levels of activity even under predation risk, thereby increasing their vulnerability (Sih *et al*. 2004). Conversely, less active individuals may forgo foraging opportunities, resulting in slower growth or reduced competitive ability. These behavioral types can create trade-offs that affect survival and resource acquisition, particularly when environmental conditions fluctuate. Such variation can influence individual fitness, shape species interactions, and alter the structure and dynamics of communities (Bolnick *et al*. 2011; McGhee *et al*. 2013).

An excellent model system for examining the effect of behavioral variation on stability of the food chain are pea aphids, *Acyrthosiphon pisum*. These aphids exhibit a well-documented and flexible set of defensive behaviors in response to predator threats, particularly dropping off the host plant. Remarkably, clonal aphids can also vary in their dropping behavior and show consistent dropping or non-dropping. This behavior is influenced by multiple factors, including predator identity, size, foraging strategy, and environmental cues. Aphids drop more frequently in response to large, active predators like adult *Coccinella septempunctata* and *Harmonia axyridis* (Losey & Denno 1998a; Francke et al. 2008), and especially when these predators are in close proximity or moving energetically (Brown 1974; Hoki et al. 2014). Alarm pheromones such as *E*-β-farnesene play a critical role in modulating dropping responses (Harrison & Preisser 2016), and predator-induced disturbance can significantly reduce aphid reproduction via non-consumptive effects (Nelson 2007).

To investigate how intraspecific behavioural variation affects food chain persistence and dynamics, we experimentally tested how the behavioral composition of prey influences these outcomes in a tritrophic system consisting of *Vicia faba* (broad bean) as the host plant resource, *Acyrthosiphon pisum* (pea aphid) as the prey, and *Coccinella septempunctata* (seven-spot ladybird) as the predator (Figure 1). Clonal aphids were screened for antipredator dropping behavior and used to establish three treatments: droppers (all individuals exhibited dropping behavior), non-droppers (none exhibited dropping behavior), and mix (a 1:1 combination of droppers and non-droppers). We tracked the persistence of the food chain over time, and used empirical dynamic modeling to estimate temporal stability and robustness of the predator-prey interaction. This approach allowed us to move beyond static or short-term snapshots and explore how intraspecific behavioral variation shapes the temporal persistence and stability of ecological interactions (Figure 1). We expected intraspecific behavioural variation in prey (antipredator dropping) to shape the temporal persistence and stability of the food chain. Specifically, we predicted that aphid populations composed exclusively of droppers (dropper treatment) would experience lower predation pressure, leading to longer food-chain persistence. In contrast, populations of non-dropping aphids (non-dropper treatment) were expected to experience higher predation pressure, resulting in shorter persistence. Mixed populations (mix treatment), containing both dropper and non-dropper aphids, were anticipated to exhibit intermediate dynamics.

**Figure 1.**
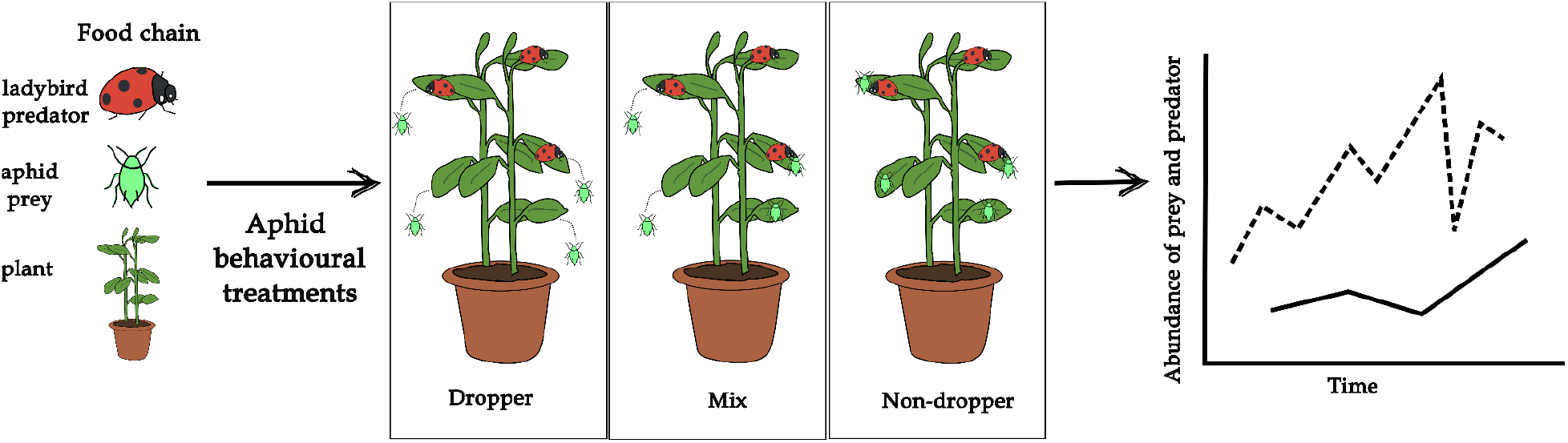
Experimental design illustrating a tri-trophic food chain composed of broad bean plants (Vicia faba), pea aphids (Acyrthosiphon pisum) as prey, and seven-spot ladybirds (Coccinella septempunctata) as predators. Clonal individual aphids were first screened for antipredator dropping behavior and classified as either ‘droppers’ if they exhibited dropping behavior or ‘non-droppers’ if they did not exhibit dropping behavior. These behaviorally distinct individuals were then used to establish three experimental treatments: dropper (all individuals drop), mix (1:1 ratio of droppers and non-droppers), and non-dropper (none drop). Each replicate consisted initially of two plants in a pot, 20 aphids, and three adult ladybirds. The setup was used to test the effect of aphid behavioral composition on aphid (prey, dashed lines) and ladybird (predator, solid line) abundance and food-chain (i.e. predator-prey) persistence (potential outcome indicated in right panel).

## 2. Materials and methods

### 2.1 Study organisms

*A. pisum* aphids were obtained from a commercial supplier (Katz Biotech, Germany) and reared on *V. faba* plants in mesh cages (60 cm x 60 cm x 60 cm, Bugdorm BD2F120; MegaView Science, Taiwan) within a climate chamber (20 °C, 16:8 h light:dark, 70% relative humidity). To establish clonal lines, single nymphs were isolated on individual plants and allowed to reproduce clonally. This process was repeated three times using nymphs acquired at different times to establish (likely) independent clonal populations. Although genetically identical, these colonies produced aphids that varied in antipredator dropping behavior (see testing below). Aphid cultures were maintained by transferring infested plant material to fresh cages when populations became denser, winged morphs appeared, or plants showed signs of wilting. All aphids used in the experiment originated from these clonal lines.

From each clonal line, young, healthy, wingless adult aphids were selected for behavioral screening. Young adults were identified by the elongated cauda, and presence of surrounding nymphs (Saberski *et al*. 2016). Individuals were gently transferred to the inverted lid of a Petri dish (5.5 cm diameter) using a moistened soft paintbrush. After placement, aphids were allowed to attach to the lid; attachment was confirmed by inverting the lid and reapplying unattached individuals. Dropping behavior was assessed by gently probing the aphid’s dorsal side of the abdomen up to three times with a modified paintbrush (all bristles removed except one) using consistent pressure. Aphids that detached and fell consistently were classified as ‘droppers’; those that remained attached were considered ‘non-droppers’. Within 2–3 hours of being classified as droppers or non-droppers, the aphids were used to set up the treatment cages.

Broad bean plants were grown from seeds (Blauetikett-Bornträger GmbH, Germany) in pots (∼25 cm tall) filled with soil (Fruhstorfer Spezialsubstrat Type T). Seeds were soaked in water for at least 24 h and seedlings were planted ∼5 cm deep in opposite corners of each pot. Plants were watered every other day and screened for use in experiments at 4–5 weeks post-germination. Only non-flowering plants with comparable height and leaf numbers (height: 40 ± 8 cm; leaf count: 51 ± 14) were selected. Plants that did not meet these criteria were either discarded or repurposed for maintaining aphid or ladybird cultures.

Adult *C. septempunctata* ladybirds were obtained from Katz Biotech (Germany) and maintained in cages with *V. faba* plants and ad libitum access to pea aphids (stock from the same supplier). Ladybird sex was determined by examining the shape of the seventh sternite (Stellwag & Losey 2014).

Experimental treatments consisted of three aphid group compositions: all droppers (dropper treatment), all non-droppers (non-dropper treatment), and a 1:1 mix of droppers and non-droppers (mix treatment), with nine replicate cages (60 cm x 60 cm x 60 cm, Bugdorm) per treatment. Each replicate cage contained 20 aphids placed on two broad bean plants growing within a single pot. After 48 h of aphid placement, three ladybirds (two females and one male) were introduced in each cage. Initial aphid and ladybird densities were informed by pilot trials that evaluated early population dynamics, ensuring aphid establishment without rapid overpopulation. Aphid (adults and nymphs) and ladybird (adults and larvae) abundances in each setup were recorded daily until day 25 after the addition of aphids, with observers blind to the cage treatment. Plant status was recorded as alive (green) or dead (brown and decaying). Note that large aphid populations could cause a plant to decline and its subsequent loss. In case of missing abundance data, we imputed values based on adjacent days. If aphid abundance was zero on both the preceding and succeeding sampling days, we imputed a value of 0 for aphids and carried forward the preceding value for ladybird abundance, based on the assumption that ladybird populations change more slowly than aphids. If aphid abundance was non-zero on both adjacent days (39 missing timepoints out of 675 total timepoints), we linearly interpolated aphid and ladybird abundances across time (using na.approx function in zoo package v1.8-13, Zeileis & Grothendieck 2005), applied separately within each cage. Plant survival, a binary variable, was forward-filled within each cage using last observation carried forward (na.locf function in zoo package v1.8-13), reflecting the assumption that once a plant is known to be alive or dead, its status persists until the next observation. In rare cases (11 instances) where aphid abundance was extremely low, aphids were recorded as absent (0) on one day but observed again the following day. Because migration was not possible in our closed system, we considered such isolated zero counts as likely false negatives due to detection error. To account for this, we substituted a minimum abundance value of 1 to reflect a detection threshold and preserve consistency in the time series. In cases where aphid abundance exceeded 2500 (in 2 of 27 cages), we capped counts at this value to maintain consistency, as higher abundances were logistically difficult to quantify accurately. In addition to above, for a subset of replicates (n = 3 per treatment), we recorded aphid abundance after the first 24 hours in the absence of ladybirds to assess aphid growth rates. Visual inspection suggested no major differences across treatments during this predator-free phase (see Figure 2, before the red dashed line). Due to logistic constraints, the experiment was conducted in multiple trials across two climate chamber. Each trial included three or six replicates, with one or two replicates per treatment.

**Figure 2.**
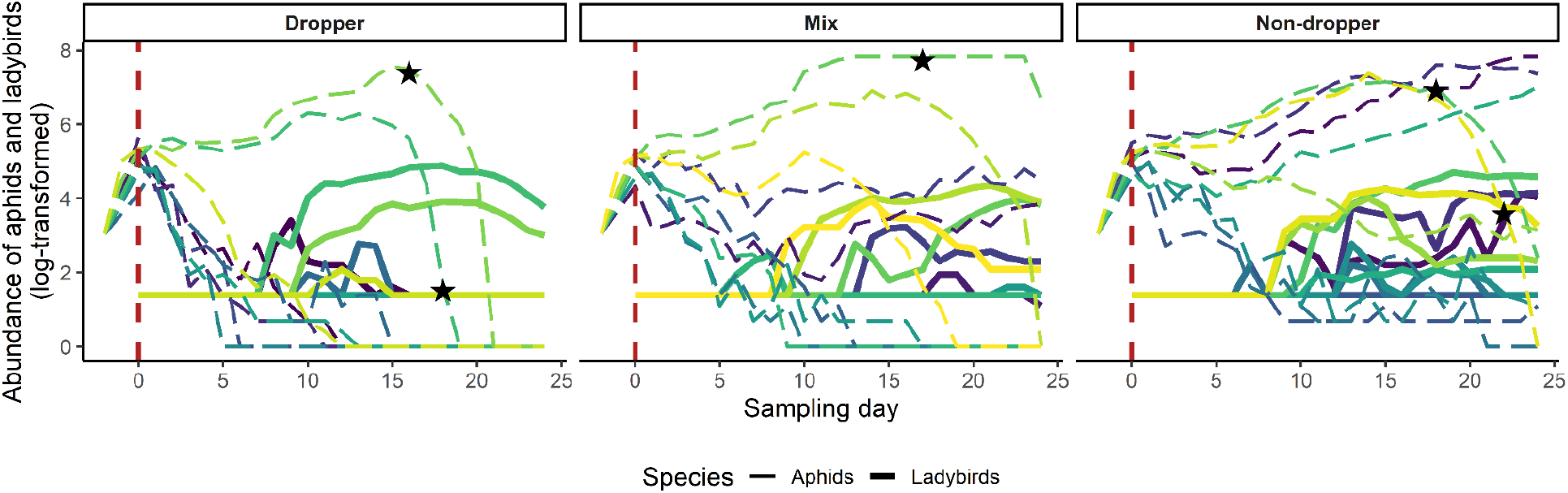
Time-series of aphid (prey) and ladybird (predator) abundances across aphid behavioral treatments: dropper (dropper aphids only), mix (both dropper and non-dropper aphids), and non-dropper (non-dropper aphids only). Log-transformed raw abundance data are shown, with a constant of one added to all values prior to transformation to account for zero values and avoid non-finite results. Dashed and solid lines represent aphid and ladybird abundances, respectively. Each colored line shows data from a single cage (n = 9 per treatment) within each treatment. The vertical red dashed line indicates the time point at which ladybirds were introduced. The black stars represent the sampling day plotted on the aphid abundance curve of the five replicate cages (two dropper, one mix and two non-dropper), where the plants died.

### 2.2 Statistical analyses of tri-trophic food-chain persistence

To assess how aphid behavioral type and within-population behavioral variation influenced the persistence of a tri-trophic food chain composed of broad beans (plant), pea aphids (prey), and seven-spot ladybirds (predator), we conducted a survival analysis using Cox proportional hazards mixed-effects models. Specifically, we tested whether prey behavioral treatments (droppers, non-droppers, or a mixture of droppers and non-droppers) impacted the time to extinction of the food chain. Extinction was defined as the first observation where any of the three trophic levels (plant, aphid, or ladybird) failed to persist within a given cage on a sampling day. For aphids and ladybirds, extinction was recorded when the observed abundance declined to zero (for aphids consistently over at least two observation points - otherwise see above). For plants, extinction was recorded when the plant was classified as dead (brown, wilted, and decaying) based on visual assessment of plant condition. Once extinction was recorded for any trophic level, the food chain for that replicate was considered extinct from that time point forward.

Survival datasets were constructed from daily repeated sampling across replicate cages. The initial model included treatment as a fixed effect, with plant age and number of leaves included as covariates to account for variation in plant condition, and stratified by species identity. To account for non-independence of observations due to the experimental structure, we included random effects for cage identity nested within aphid clone, and for the position of the cage nested within the climate chamber. Mixed-effects cox proportional hazards models were fitted using the *coxme* package (version 2.2.22, Therneau 2024a) using the survival package to construct survival objects (version 3.8.3, Therneau 2024b). Model selection was conducted based on the Akaike Information Criterion corrected for small sample sizes (AICc), by comparing full and reduced models (see Supplementary Table S1). Hazard ratios with corresponding standard errors were extracted from the best-fitting model, which included treatment as a fixed effect and cage ID nested within aphid clone as a random effect. In addition to the overall food chain persistence analysis, we also generated species-specific survival curves for plants, aphids, and ladybirds to visualize how behavioral treatments influenced persistence at each trophic level. For these, separate survival models were fitted for each species, and combined survival curves were generated by mapping treatment groups as color and species identity as linetype.

To verify the robustness of our results, we also performed survival analyses excluding plant mortality and defining extinction based solely on aphid and ladybird persistence, for which abundance data was available. This analysis yielded qualitatively similar treatment effects on food chain persistence (see Supplementary Figure S1). All statistical analyses were conducted using R (version 4.4.2).

### 2.3 Statistical analyses of state transitions observed

We evaluated how aphid behavioral type and within-population behavioral variation influenced transitions between states over time in the tri-trophic food chain. In theory, seven states could arise from the presence/absence combinations of plant, aphid, and ladybird in our experimental food-chain: (1) plant–aphid–ladybird (full food chain), (2) plant–aphid, (3) plant–ladybird, (4) aphid–ladybird, (5) plant only, (6) aphid only, and (7) ladybird only. However, in our experiment, we primarily observed four states: (1) plant–aphid–ladybird present (full food chain), plant–ladybird (aphid extinct), aphid–ladybird (plant extinct), and ladybird only (both plant and aphid extinct). The food chain transitions were primarily observed from the full food chain (plant–aphid–ladybird) to either plant–ladybird or aphid–ladybird states, followed by further simplification to ladybird-only (Supplementary Table S2). To quantify how aphid behavioral treatments influenced transitions from the full food chain to simpler states, we fitted a multi-state Markov model, where transitions occurred upon the absence of one or more species (msm package version 1.8.2, Jackson 2011). Hazard ratios (HRs) with 95% confidence intervals were estimated to evaluate treatment effects on these transitions. Specifically, we tested the transitions from the full food chain to the simplified states plant–ladybird and aphid–ladybird. In addition, empirical state occupancy proportions over time were calculated directly from observed cage-level state frequencies and plotted.

### 2.4 Statistical analyses of dynamical stability of the tri-trophic food chain (prey-predator abundance)

We evaluated whether aphid behavioral types and within-population behavioral variation impacted the dynamical stability of the tri-trophic food chain over time. To do this, we used empirical dynamical modelling (EDM) (Deyle *et al*. 2016; Sugihara *et al*. 2011). EDM extends classical community matrix calculation which is defined for ecological systems at equilibrium. The community matrix is defined as the matrix that encapsulates the behavior of the system at equilibrium, which mathematically contains the partial derivatives of the system, i.e., how the per-capita growth of a species is impacted when there is a change in the abundance of other species at equilibrium. However, ecological systems are rarely at equilibrium, and the strength of species interactions can change non-linearly over time. EDM then quantifies how such interaction strengths change over time to provide a dynamical perspective of the stability of the system. EDM reconstructs the underlying dynamical system using observed time series by an equation-free mechanistic modelling approach. Briefly, the state of the food chain is defined by the aphid abundance and ladybird abundance. This state maps out a trajectory of the system in a state-space from which strengths of species interactions can be estimated, by a locally weighted multivariate linear regression (Deyle *et al*. 2016). These interaction strengths are then used to construct the Jacobian matrix from which local dynamical stability is estimated. To elaborate on the methodology, the state-space is first constructed by plotting the aphid-ladybird time series against each other. Thus instead of aphid abundance over time, or ladybird abundance over time, aphid and ladybird time series are plotted on the two axes (for details see Deyle *et al*. 2016). Now, each point in the state space is a time point that characterizes the x-y state coordinate of the system. Next, by using locally weighted multivariate linear regression for each time point, we approximate the best model by giving weights to the points in the state-space that are near to the point where the estimation is taking place. This is done sequentially over each time point. With that, estimates of Jacobian elements of our 2×2 system are calculated where the diagonal of the Jacobian matrix captures the intraspecific strengths, and off-diagonal captures inter-specific strength. From such estimation of the Jacobian matrix, one can then directly estimate the dominant eigenvalue, which is the local stability of the system. Next, the average robustness of the system is estimated as a geometric mean of all the eigenvalues over time. Average robustness was calculated once the Jacobian matrix was estimated from the time series data of pea aphids and ladybird time series using S-map from EDM. We thus evaluated how the local stability and average robustness were impacted by the behavioral treatment types of the model. However, while estimating average robustness and local stability, we had to exclude seven (four dropper, two mix and one non-dropper cage) out of twenty-seven cages as they did not fit the data resolution criteria for S-map analyses. The data resolution criteria for such analyses need some amount of fluctuation or changes in abundances of one of the state variables. In our case, in the above mentioned six cages, there were no changes or fluctuations observed in the ladybird abundances over time.

## 3. Results

### 3.1 Food-chain persistence

Overall, raw abundance data showed that aphid populations declined sharply in the dropper treatment (i.e., where all aphids exhibited dropping behavior), but remained more variable in the mix and non-dropper treatments (Figure 2). Ladybird abundance declined most rapidly in the dropper treatment and least in the non-dropper treatment, although they did not go extinct in any replicate. In two dropper, one mix and two non-dropper replicate cages, the plants also collapsed during the experiment period.

Aphid behavioral type (dropper or non-dropper) and behavioral variation (no variation: all droppers or all non-droppers; variation: a mix of both types) influenced food chain persistence (Figure 3A). The Cox proportional hazards mixed-effects model showed that the extinction rate ratio (i.e., the exponentiated hazard estimate) was highest in the dropper treatment, which served as the reference level (Figure 3B). In comparison, the extinction rate ratio was significantly lower in the mix treatment (β = –0.94 ± 0.44 SE, *p* = 0.033; hazard ratio [HR] = 0.39) and even further reduced in the non-dropper treatment (β = –2.59 ± 0.51 SE, *p* < 0.001; HR = 0.07). Thus, the dropping behavior negatively affected the persistence of the food chain. When stratified by species, aphid persistence declined most rapidly in the dropper treatment and was highest in the non-dropper treatment, mirroring the overall food chain persistence pattern (Figure 3C). Ladybird persistence remained high across all treatments, as ladybird extinction (zero abundance) was rarely observed, although some gradual declines in abundance occurred over time. Plant survival did not differ substantially across treatments.

**Figure 3.**
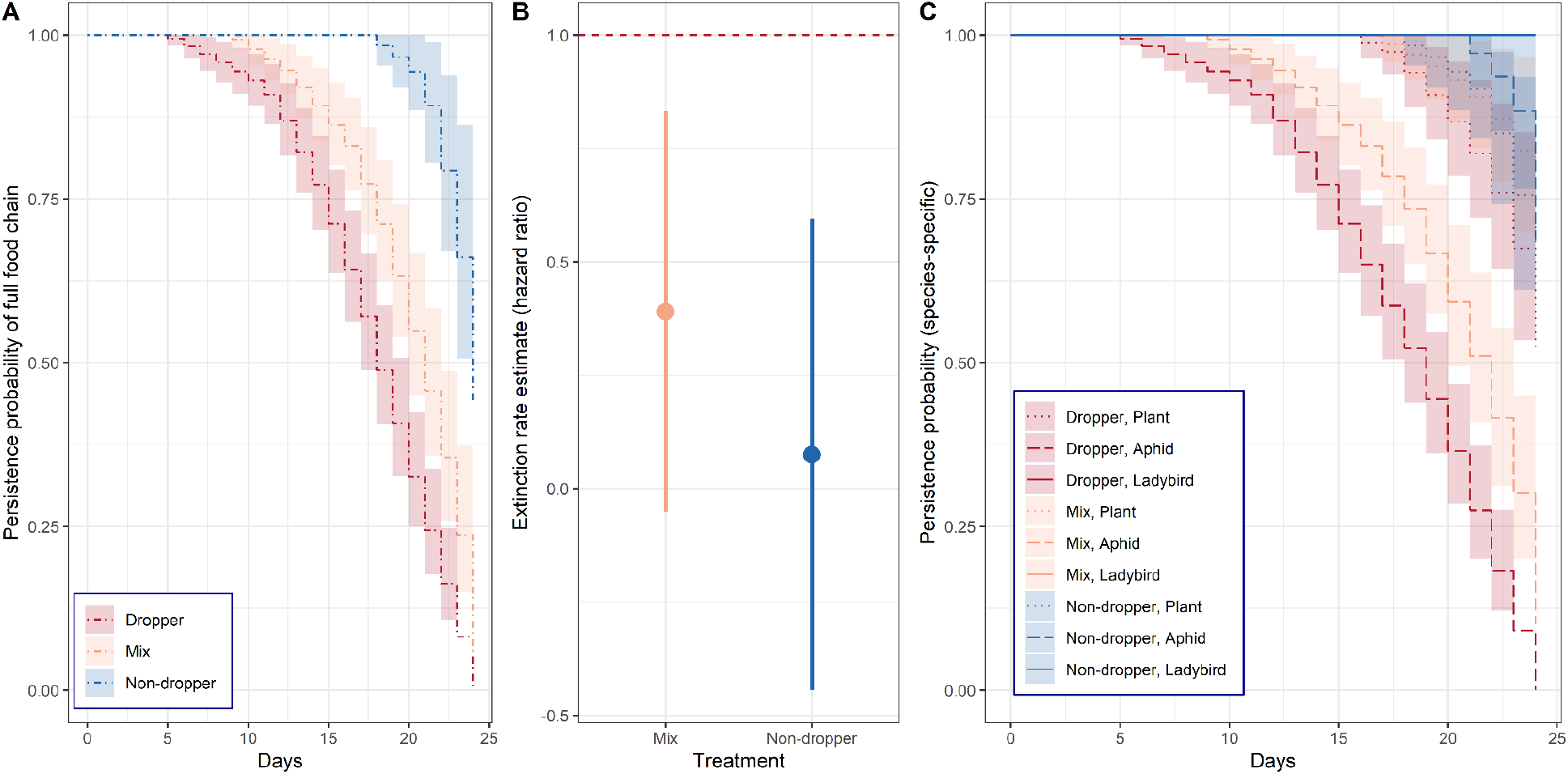
Persistence probability of the tri-trophic food chain across the three aphid behavioral treatments (dropper, mix, and non-dropper): (A) Cox survival curves showing overall food chain persistence probability under each behavioral treatment. (B) Cox model estimates of extinction risk (hazard ratios ± SE) for each treatment relative to the dropper treatment (reference, dashed red line). Both mix and non-dropper treatments exhibited significantly lower extinction risk compared to the dropper treatment. (C) Species-specific survival curves showing persistence probability separately for plants, aphids, and ladybirds under each treatment. Aphid persistence declined most rapidly in the dropper treatment, while ladybirds and plants generally showed higher persistence across treatments.

### 3.2 Tri-trophic food chain transitions

Multi-state Markov models revealed that aphid behavioral treatments significantly affected transition rates. The hazard of aphid extinction (transition from full food chain to plant–ladybird) was substantially reduced under the non-dropper treatment relative to dropper (HR = 0.07; 95% CI: 0.01–0.53), indicating a ∼93% lower risk of aphid loss. The mix treatment also reduced the aphid extinction hazard relative to dropper (HR = 0.54; 95% CI: 0.18–1.61), although this effect was not statistically significant. In contrast, transitions from full food chain to aphid–ladybird state (plant extinction) were not strongly influenced by treatment. Hazard ratio estimates for these transitions were lower than 1 for both mix (HR = 0.32; 95% CI: 0.03–3.48) and non-dropper treatments (HR = 0.46; 95% CI: 0.06–3.27), but confidence intervals overlapped unity. Empirical state occupancy patterns over time supported these effects (Figure 4), with the full food chain persisting longest in the non-dropper treatment and simplified states emerging earliest under dropper.

**Figure 4.**
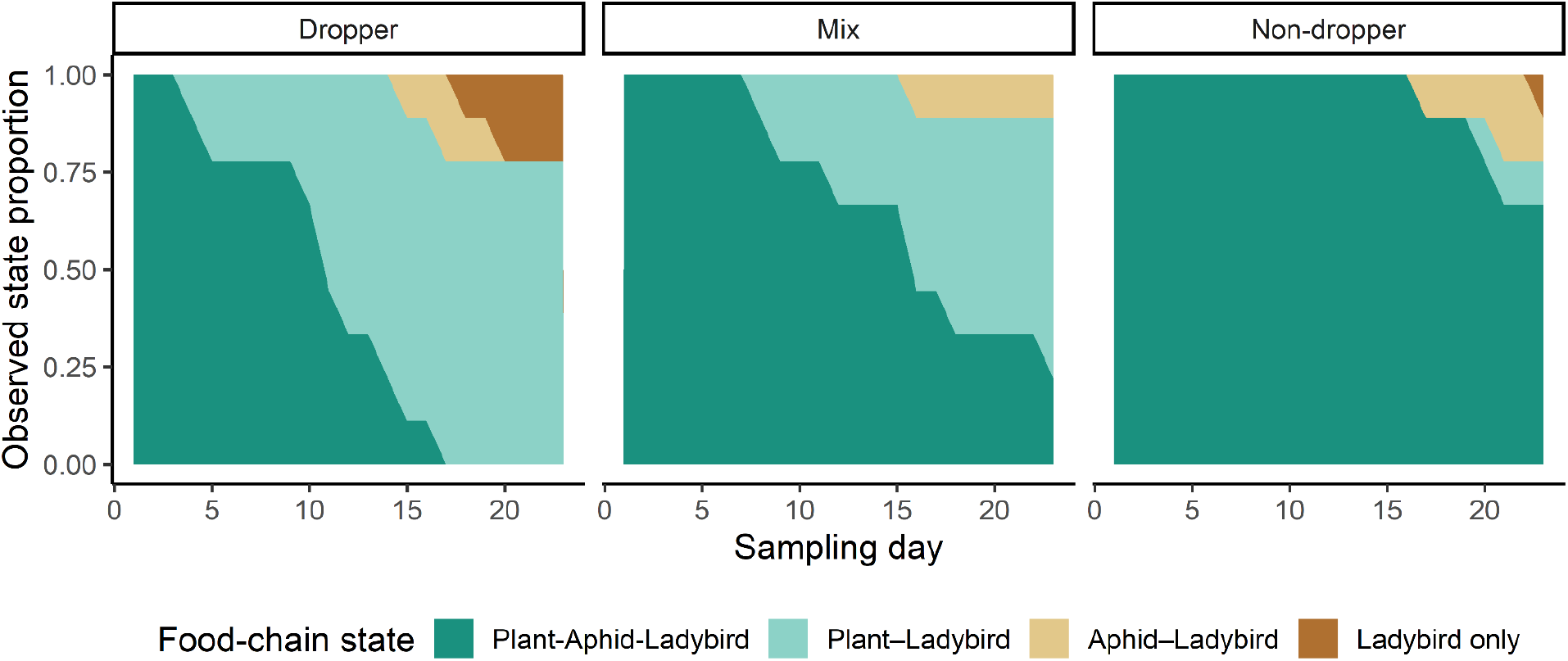
Empirical occupancy dynamics of tri-trophic food chain states across aphid behavioral treatments (dropper, mix, non-dropper). Shaded areas represent the observed proportion of cages occupying each state at each sampling day. States are defined by species presence: full food chain (plant–aphid–ladybird), plant–ladybird (aphid extinct), aphid–ladybird (plant extinct), and ladybird only (both plant and aphid extinct). The tri-trophic structure (dark green) progressively simplified as species were lost, with dynamics varying by treatment: aphid extinction occurred earlier in dropper and mix treatments, while non-dropper cages retained the full food chain for longer.

### 3.3 Dynamical stability of the tri-trophic system

Our results indicated that all food chains were generally unstable (in the beginning) indicated by positive values of the dominant eigenvalue in general (Figure 5A). While dropper treatments showed a trend towards increasingly negative eigenvalues over time indicating more stability, this pattern was accompanied by high variability, and confidence intervals overlapped with those of the non-dropper and mix treatments. A similar trend was observed for average robustness (Figure 5B): dropper systems appeared to increase in robustness during later sampling days, while non-dropper and mix treatments remained relatively less robust over time. However, overlapping confidence intervals again preclude strong inferences about treatment-level differences. For individual cage dominant eigenvalues (Re(λ)) and robustness, see Supplementary Figures S2, S3. These findings indicated that while the non-dropper behavior of the aphids might facilitate longer-term persistence of the entire food chain, it was not associated with local stability or robustness. Taken together, our results suggested possible behavioral modulation of dynamical properties, but further replication would be needed to robustly assess treatment effects on dynamical food chain stability.

**Figure 5.**
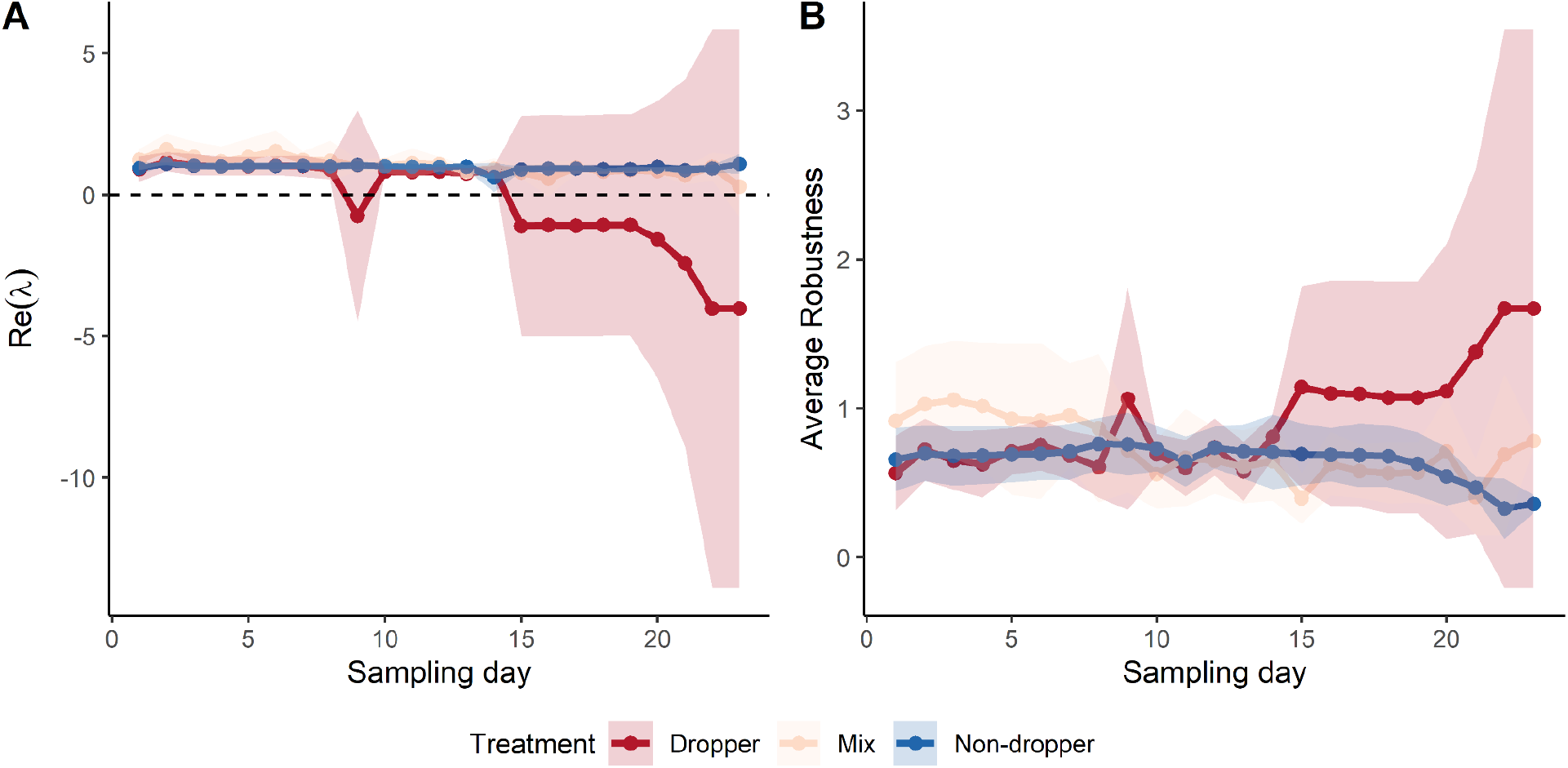
Local stability and average robustness of the tri-trophic food chain (prey-predator abundance) over time under different aphid behavioral treatments. (A) Dominant eigenvalue (Re(λ)) and (B) average robustness estimated from S-map analysis using empirical dynamic modeling, based on aphid and ladybird abundance time series. Negative Re(λ) values indicate local stability, while average robustness represents the geometric mean of local Jacobian eigenvalues over time. Higher values indicate greater robustness. Lines represent the mean across replicates; shaded regions indicate 95% confidence intervals. Although the dropper treatment showed a trend toward greater stability and robustness during later stages of the experiment, confidence intervals overlapped across treatments.

## 4. Discussion

Behavioral variation among individuals is typically attributed to underlying genetic differences or environmental influences (Dochtermann & Royauté 2019; Laskowski *et al*. 2022). However, emerging evidence suggests that additional factors may also contribute to such variation (Laskowski *et al*. 2022; Sih *et al*. 2004). While trait variation linked to genotypic differences has been shown to influence community diversity (Banitz 2019; Barabas & D’Andrea 2016; Baruah *et al*. 2025), composition, stability, and population persistence (Arroyo-Correa *et al*. 2023; Barbour *et al*. 2016, 2019; Baruah 2022; Gibert 2016; Hogle *et al*. 2022), the ecological consequences of behavioral variation, particularly its effects on biodiversity, stability, and community persistence, remain relatively unexplored, especially in experimental contexts. Quantifying how behavior shapes ecological stability could offer valuable insights into how individual-level differences scale up to affect broader community and ecosystem processes. Dropping behaviour, observed in pea aphids, is a common antipredator strategy, wherein individuals detach from their host plant in response to disturbance cues such as predation risk, providing an immediate survival advantage (Humphreys & Ruxton 2019). Numerous studies have examined the biotic and abiotic conditions that trigger this behavior and its ecological consequences (Agabiti *et al*. 2016; Braendle & Weisser 2001; Gish *et al*. 2012; Harrison & Preisser 2016; Humphreys *et al*. 2021; Nelson 2007; Schuett *et al*. 2015). However, it remains unclear whether such behavioral variation can scale up to affect the persistence of a simple food-chain community.

Recent work had emphasized that in natural settings, aphid populations might be composed entirely of clonal individuals. This suggests that the use of clonal lines in experimental research on aphid behavior is not only appropriate but ecologically realistic (Bühler & Schweiger 2024). In line with this notion, studies have documented behavioral variation among aphid clones, even under controlled conditions (Schuett *et al*. 2011). For example, previous studies have observed consistent differences in escape responses among red and green pea aphid clones (Braendle & Weisser 2001; Schuett *et al*. 2015). Red clones showed a greater tendency to drop under artificial threats, although this trend did not always hold in natural predator scenarios (Braendle & Weisser 2001). Such inconsistencies point to the challenge of extrapolating laboratory-based behavioral assays to field conditions and emphasize the complexity of interpreting behavioral plasticity in natural ecological settings. Aphid responses to predators are shaped by a combination of prey as well as predator characteristics and the environmental context. With regard to predator characteristics, their identity, size, and foraging behavior significantly influence the likelihood of aphid dropping. For instance, larger and more active predators such as adult *Coccinella septempunctata* and *Harmonia axyridis* prompt significantly more frequent dropping responses than smaller or less active predator species (Francke *et al*. 2008; Losey & Denno 1999). Francke et al. (2008) further demonstrated that older and more active *H. axyridis* individuals induced more pronounced dropping responses compared to younger conspecifics. Similarly, earlier studies have indicated that close proximity and high predator movement intensity are critical triggers (Brodsky & Barlow 1986; Brown 1974; Hoki *et al*. 2014). The effects of predator activity may be mediated through multiple sensory pathways, including vibrational cues transmitted via the plant (Clegg & Barlow 1982; Montgomery & Nault 1977). Additionally, predators can become coated with aphid alarm pheromones, such as *E*-β-farnesene, thereby amplifying perceived predation risk and increasing the likelihood of dropping (Mondor & Roitberg 2004).

Our study demonstrated that intraspecific variation in prey behavioral strategy could shape the persistence of tri-trophic food chain. Specifically, we found that aphid dropping behavior, while known to reduce immediate predation risk, significantly decreased food-chain persistence (dropper treatment). In contrast, both the mix and non-dropper treatments exhibited longer persistence (Figure 2). Notably, the non-dropper treatment showed the highest persistence overall, suggesting that avoiding dropping behaviours might enhance the long-term stability of the entire food chain under sustained ecological interactions. This was in contrast to our prediction where we expected dropper aphids to experience lower predation pressure and a prolonged food-chain persistence. This unexpected result suggested that the benefits of dropping behavior (reducing predation risk) might be outweighed by energetic or demographic costs, ultimately reducing long-term persistence. Indeed, while dropping might provide an effective means of evading predators (Francke *et al*. 2008; Losey & Denno 1999), it also incurred energetic and time-based trade-offs. In pea aphids, repeated dropping behaviour delayd development and reduces reproductive output, highlighting substantial non-consumptive costs that could constrain population growth (Agabiti *et al*. 2016). Moreover, frequent dropping could increase vulnerability to ground-based predators (absent in our study) and could expose aphids to suboptimal microhabitats, increasing the ecological costs of such a defence strategy (Nelson 2007). Additionally, dropping was suggested to be employed selectively under high-risk conditions, given such high energetic cost and substantial fitness consequences (Harrison & Preisser 2016). Indeed, non-consumptive effects such as reduced feeding and stress-induced reproductive suppression could be as ecologically influential as direct predation, and models excluding those effects might overestimate population growth by up to 35% (Fill et al. 2012; Nelson et al. 2004; Nelson 2007). Thus, our observed lower persistence of aphids in the dropper treatment was consistent with such physiological and fitness costs accumulating over time, although we did not explicitly measure temporal fitness of the droppers. We also observed a trend of higher ladybird abundances in more non-dropper replicates compared to dropper ones (Figure 2), suggesting that reduced aphid persistence might cascade upwards to affect predator dynamics. Larger and more active predators such as *C. septempunctata* were known to elicit stronger dropping responses from aphids (Francke et al. 2008; Hoki et al. 2014), but when prey frequently escapes, the predator foraging efficiency might decline. Thus, the lower predator abundance in dropper treatments might reflect reduced prey availability and increased handling or search time under high prey evasiveness.

Interestingly, although mix treatment food chains persisted longer than those composed solely of droppers, they did not exceed the persistence observed in non-dropper treatments (Figures 2–3), suggesting that behavioural diversity may buffer, but not fully offset, the associated costs of dropping behaviour. Similarly, the mix treatment also maintained a higher frequency of the tri-trophic system state (full food chain) over time than the dropper treatment. This suggested that within-population behavioral variation might stabilize the food chain by reducing the overall intensity of non-consumptive costs at the population level of the prey. In comparison, the dropper treatment frequently transitioned into simplified food chain states (e.g., plant-ladybird), highlighting the destabilizing impact of such an antipredator strategy that carries high energetic costs. The buffering effects of trait variation have been predicted in theoretical models (Barabás *et al*. 2022), but in these models trait variation has usually been modelled to be genetic in nature. Such variation in interaction strength arising due to variation in species has been shown to be beneficial in promoting stability and feasibility in mutualistic communities (Baruah 2022, Baruah & Wittmann 2024), but species coexistence is disrupted in competitive communities (Barabas & D’Andrea 2016; Baruah *et al*. 2025; Hart *et al*. 2016). In our experiment, clonal aphids were used that nevertheless showed variation in dropping behavior. The impacts of such behavioral heterogeneity on predator-prey persistence and stability remain rarely tested in empirical systems.

Our empirical dynamic modeling revealed treatment-specific trends in the temporal stability and robustness of the tri-trophic food chain. While systems with dropper aphids exhibited an apparent increase in local stability (more negative Re(λ)) and robustness over time, these trends were accompanied by high variability, and confidence intervals overlapped across treatments. Thus, while the patterns were suggestive, they should be interpreted with caution. Towards the end of the experiment, predator–prey dynamics in the dropper treatment appeared more stable (as reflected in dominant eigenvalue estimates), despite lower overall persistence. This likely resulted from the extinction of aphids in several replicates, after which predator abundances remained fixed at three individuals through day 25. In these cases, the dynamics effectively reduced to a single, non-fluctuating predator population (a fixed point), artificially stabilizing the system. However, under natural conditions, predator extinction would likely follow due to lack of prey to feed upon. Such trade-offs between ecological persistence and dynamical stability are rarely empirically quantified and highlight the importance of integrating behavioral traits into food-web models.

In conclusion, our findings contribute to a growing body of evidence indicating that behavioral variation has complex, community and system-level consequences. While dropping behavior provides a clear immediate benefit for the prey, its chronic use imposes significant costs that can reduce prey and predator persistence and restructure trophic interactions. Our study extends this understanding by showing that behavioral variation within prey populations can buffer these effects and promote food chain stability. These insights emphasize the importance of considering non-consumptive effects and trait variation in studies of community dynamics and stability, especially in systems where behavior mediates ecological feedback.

## Data Availability

The dataset will be uploaded to Zenodo.

## Acknowledgments

GB acknowledges Deutsche Forschungsgemeinschaft (DFG) Walter Benjamin grant number BA 7974/1-1. GB and PS acknowledge the Faculty of Biology for a small experimental Grant supporting the performance of the experiment. We thank Corvin Alpmann for assistance with the experiment, Rabea Schweiger for helpful discussions, and Christine Fiebig and Karin Djendouci for logistical support.

