## appendix for "Behavioral variation affects persistence of an experimental food-chain"

### Supplementary Information

Supplementary Table S1. Model selection results for the food chain persistence Cox proportional hazards mixed-effects models. We compared models using Akaike Information Criterion corrected for small sample size (AICc). The best-fitting model (highlighted in bold) retained Treatment as a fixed effect and included random effects for Cage ID nested within Aphid clone.

| Model | Fixed effects | Random effects | K | AICc | ΔAICc | AICc weight |
|---|---|---|---|---|---|---|
| full | Treatment + Plant_Age + No_of_leaves | (Climate chamber/Position of cage) + (Aphid clone/Cage ID) | 8 | 1785.52 | 3.54 | 0.1 |

| | | | | | | |
|---|---|---|---|---|---|---|
| reduced 1 | Treatment + Plant_Age | (Climate chamber/Position of cage) + (Aphid clone/Cage ID) | 7 | 1783.81 | 1.83 | 0.23 |
| reduced 2 | Treatment | (Climate chamber/Position of cage) + (Aphid clone/Cage ID) | 6 | 1785.41 | 3.43 | 0.1 |
| **reduced 3** | **Treatment** | **(Aphid clone/Cage ID)** | **4** | **1781.98** | **0** | **0.57** |

K = number of parameters; ΔAICc = difference from best model; AICc weight = relative model support.

Supplementary Table S2. Observed counts of state transitions among food-chain states during the experiment. Rows indicate the state at time t, and columns indicate the state at time t+1.

| From/to | Ladybird only | Plant-ladybird | Aphid–ladybird | Plant–aphid–ladybird |
|---|---|---|---|---|
| Ladybird only | 8 | 0 | 0 | 0 |
| Plant-ladybird | 0 | 149 | 0 | 0 |
| Aphid–ladybird | 3 | 0 | 18 | 0 |

| | | | | |
|---|---|---|---|---|
| Plant–aphid–ladybird | 0 | 14 | 5 | 397 |

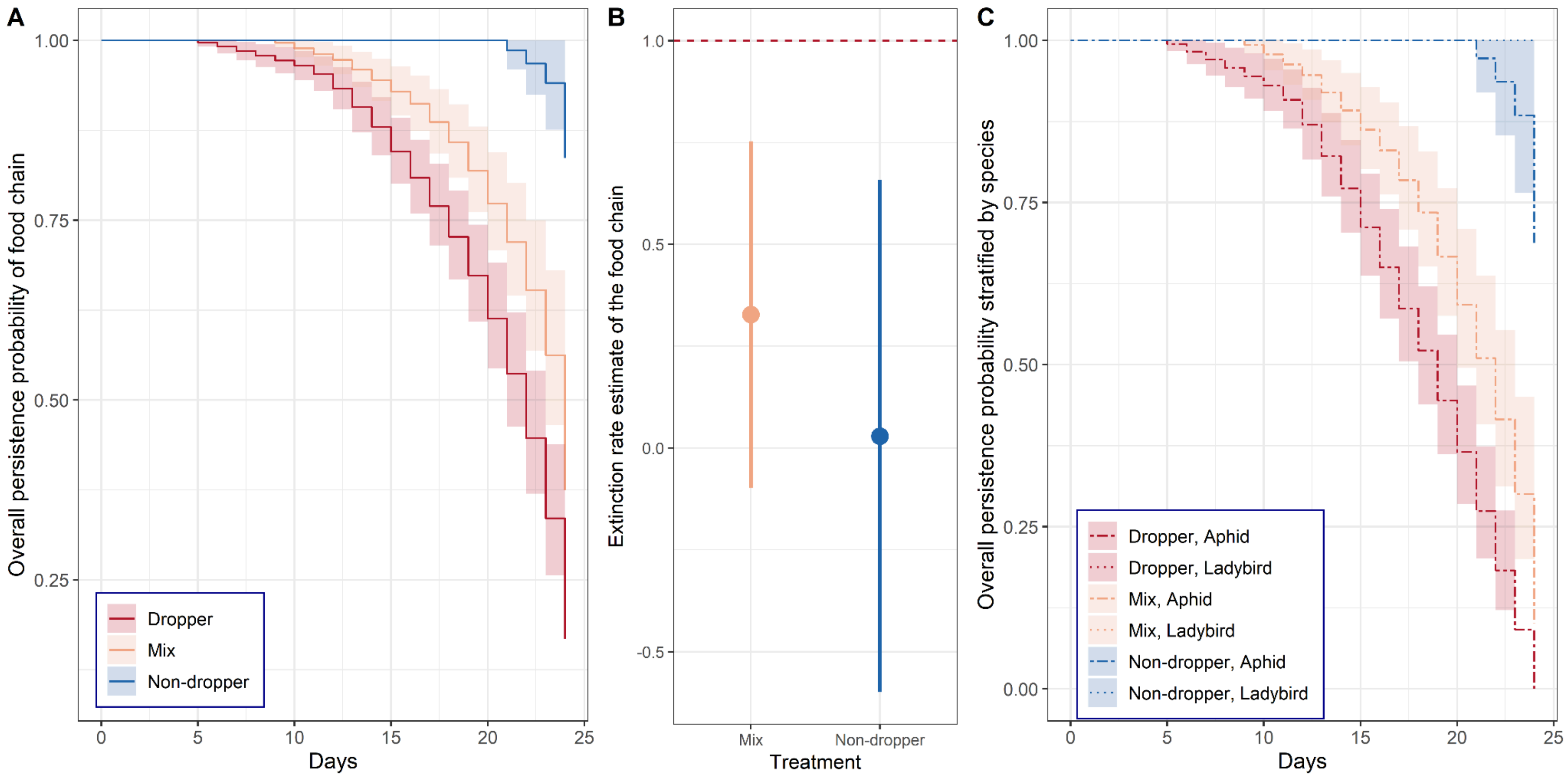


Supplementary Figure S1. Food chain persistence when extinction was defined only by aphid and ladybird persistence (plant survival not included). (A) Survival curves showing overall food chain persistence probability across the three behavioral treatments. (B) Hazard ratio estimates (± SE) for treatment effects on extinction rate, using the dropper treatment as reference. (C) Species-specific persistence curves for aphids and ladybirds across treatments. Results are qualitatively similar to the main analysis that included plant mortality (see Figure 3).

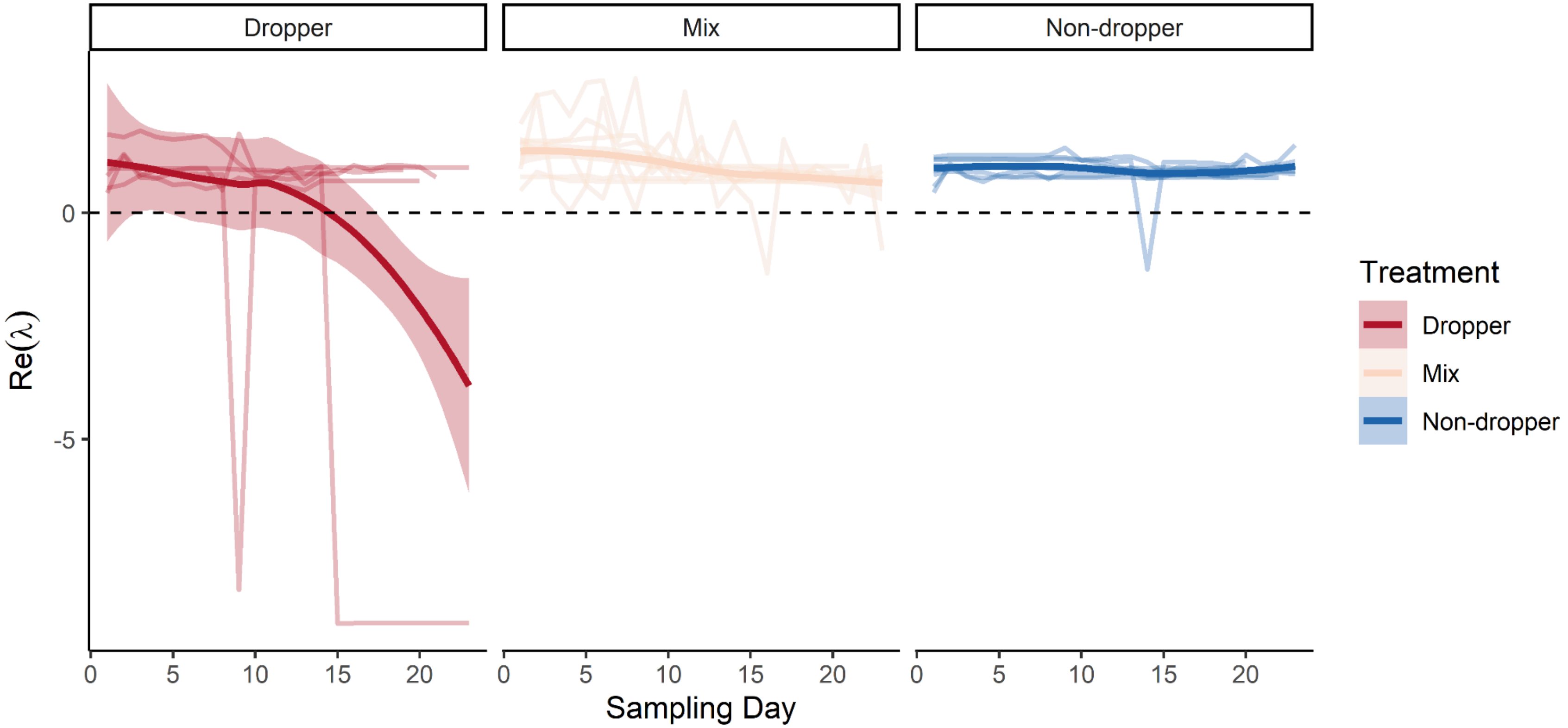


Supplementary Figure S2. Dominant eigenvalue trajectories of individual cages across aphid behavioral treatments. Each line represents the estimated dominant eigenvalue (Re(λ)) over time for a single cage. Lighter lines show raw trajectories; bold lines represent LOESS fits with 95% confidence intervals for each treatment. While average trends suggest negative Re(λ) values in dropper treatments over time, substantial variation is present among replicates. Dashed horizontal line at Re(λ) = 0 denotes the local stability threshold.

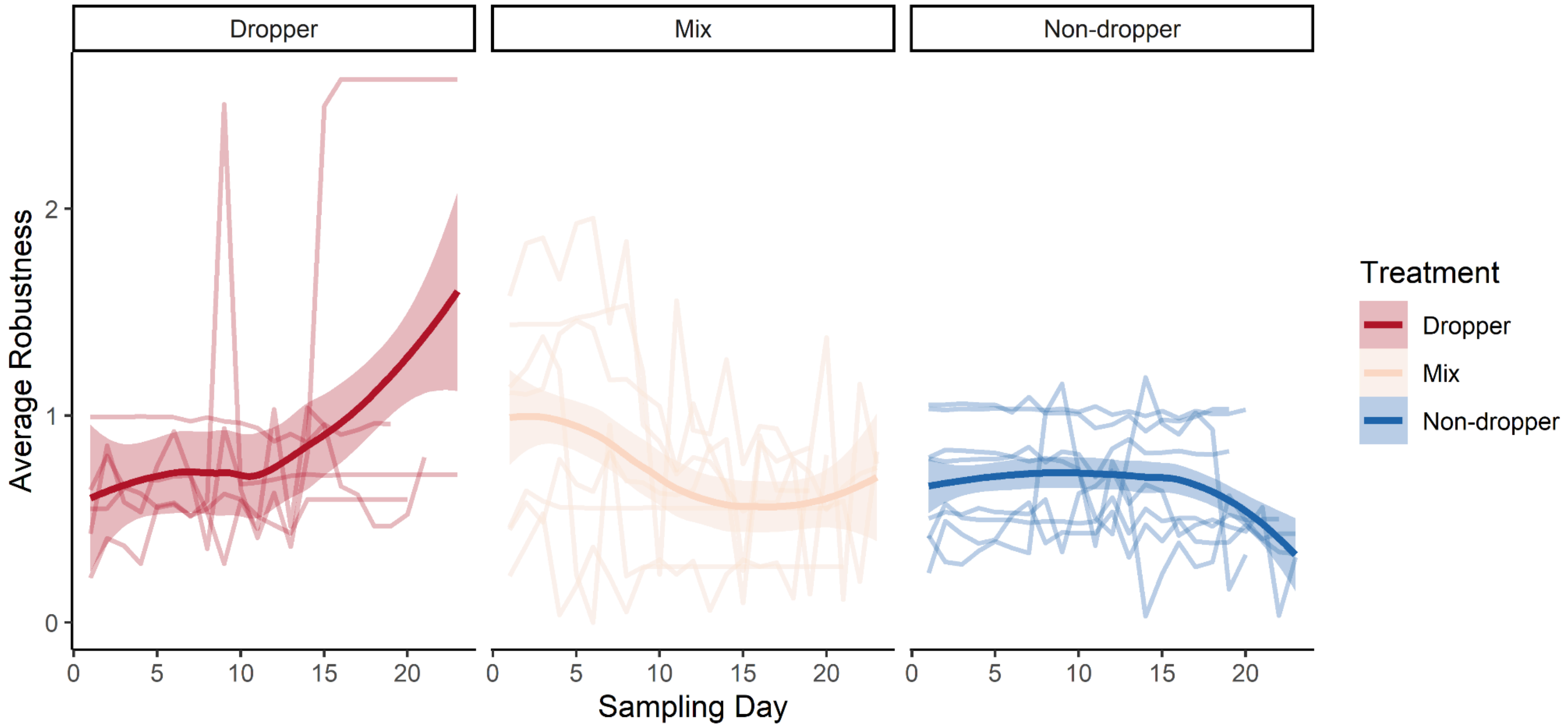


Supplementary Figure S3. Individual cage-level dynamics of robustness across aphid behavioral treatments. Each line represents the estimated average robustness (geometric mean of local Jacobian eigenvalues) over time for a single cage. Solid curves show LOESS fits with shaded 95% confidence intervals for each treatment. While robustness appeared to increase in the dropper treatment over time, substantial variability was observed across cages, and confidence intervals overlapped among treatments.
